# Primer Blind Spots: Underrepresentation of ESKAPE Pathogens from Developing Regions in 16S rRNA Targeted Sequencing

**DOI:** 10.1101/2025.11.17.688800

**Authors:** Divya Dilip Gawande, Khushi Chouhan, Ishaan Gupta

## Abstract

Targeted 16S rRNA sequencing is stalwart of microbial profiling and diagnostics, but its reliance on some “universal” primers introduces bias. In this in-silico study, we investigated how well these primers capture ESKAPE pathogens by screening multiple databases and found that a considerable fraction showed suboptimal primer binding, suggesting that amplification and identification may be compromised. Notably, these strains were disproportionately associated with isolates from developing countries, underscoring the risk of primer bias in global surveillance.

## Introduction

Antimicrobial resistance (AMR) is a pressing global health threat, driven by diagnostic delays and inadequate surveillance of pathogens (1). Targeted 16S rRNA gene sequencing is used for rapid pathogen identification; however, its accuracy can be affected by various factors, including primer selection, sequencing platforms, assembly protocols, quality control parameters, and clustering methods (2–4). A critical gap exists where a significant portion of known bacterial strains, particularly those from underrepresented geographical regions, may not be adequately captured by these primers, leading to a silent epidemic of undiagnosed and misidentified pathogens(5). Strains from developing countries, where sequencing capacity is often limited, may be particularly vulnerable to underrepresentation if standard primer sets fail to amplify them(6,7).

The variability observed in bacterial profiling highlights the need for standardized methodologies to ensure consistency(8) by employing multiple primer sets and updated, curated databases to enhance taxonomic resolution and reduce bias(9,10).

Here, we present a systematic in-silico analysis of 16S rRNA sequences from ESKAPE pathogens using multiple public repositories. We assess the ability of 16S primer binding, quantifying isolates lacking primer sites, and evaluated whether primer bias disproportionately affects strains from low-income regions. Our findings highlight the need for more inclusive primer strategies to strengthen global diagnostic and surveillance efforts.

## Materials and Methods

The 16S rRNA gene sequences (greater than 1000 bp) of six ESKAPE bacterial species were retrieved from GSR (n = 9538), MIMt (n = 1323), and NCBI (n = 793). Universal primer sequences complementary to conserved regions flanking hypervariable sequences were extracted from previously published literature (11) were then employed for systematic searches across relevant databases.

Sequence parsing, primer alignment, coverage analysis, and result visualization were carried out using a combination of Bash and Python scripts (version 3.9). Input FASTA files were processed alongside a primer dictionary, and each sequence was searched for forward and reverse primers by aligning the forward primer to the original strand and the reverse primer to the reverse complement. Exact binding positions were identified using regular expression searches, and amplicon lengths were calculated from primer coordinates. To account for primer bias, we implemented an additional pipeline allowing up to two mismatches, rejecting mismatches at the 3′ end but permitting them at internal and 5′ positions. This was achieved using a sliding-window approach with multiprocessing for efficiency, and mismatch details were recorded for each primer pair. A secondary analysis quantified the proportion of sequences containing forward, reverse, or both primers and calculated mean amplicon lengths across species.

To evaluate whether multiplexing primers improved detection, we generated a coverage matrix of species versus primer presence. A greedy algorithm was applied to iteratively select the primer pair that maximized coverage of previously uncovered species at each step. Coverage statistics were recorded at each step, and plots were generated in Matplotlib.

## Metadata Retrieval and Geographical Mapping

Accession identifiers corresponding to sequences lacking primer matches were used to retrieve sample metadata from NCBI. The pipeline employed Biopython’s Entrez module to query the E-utilities API, processing accession IDs in parallel batches with up to five threads via ThreadPoolExecutor. Records were fetched in GenBank XML. For each valid accession, metadata fields including accession number, organism, strain, sequence type, sequence length, geographical origin, isolation source, and associated references were extracted and compiled into Excel files. To ensure robustness, the code retried each accession in different formats up to three times.

## Results

To establish the genomic context of widely used universal primers, we extracted primer sequences from published literature and mapped them against the *E. coli* 16S rRNA gene. Primer pairs such as 283 (targeting V3–V4), V3, and V4.1 overlap significantly with hypervariable regions, enabling broad taxonomic resolution. In contrast, others, such as V2 and V7, capture distinct loci with narrower coverage potential.

Analysis of primer binding across these databases (n = 3, x = 11654) revealed that **primer pair 283 (V3–V4)** was the most frequent, present in **97.73% of species**, followed by **V3 (90.92%), V4.1 (55.74%)**, and **V5.1 (52.46%)**. The most common amplicon sizes were **282 bp (283), 188 bp (V3), 152 bp (V4.3)**, and **149 bp (V5.2)**. At the genus level, the highest match rates were observed in **Pseudomonas (47.79%), followed by Klebsiella, Enterobacter, and Staphylococcus (15.76%)**, suggesting that primers 283 and V3 are particularly effective for ESKAPE pathogens. By contrast, primers **V4.3, V5.2, and V6** mapped to fewer species. When mismatches were permitted (up to 2–3 bases, excluding 3′ end mismatches), coverage of **V4.2, V6, and V7** primers increased above **60%** across all three databases, demonstrating mismatch tolerance increases primer utility for broader species detection.

To determine optimal primer combinations, a greedy algorithm was applied to cumulative coverage (see *Figure Z*, second graph). **Primer 283** provided the strongest initial coverage. Adding **V6** significantly improved coverage, and incorporating V3 increased detection to over **90% across all databases**. Beyond this, additional primers such as **V5.1** and **V4.1** only marginally increased coverage, indicating a plateau. The combination of **283, V6, and V3** thus represents the most effective set for comprehensive detection of ESKAPE and other clinically relevant species(Figure 1).

**Figure 1:**
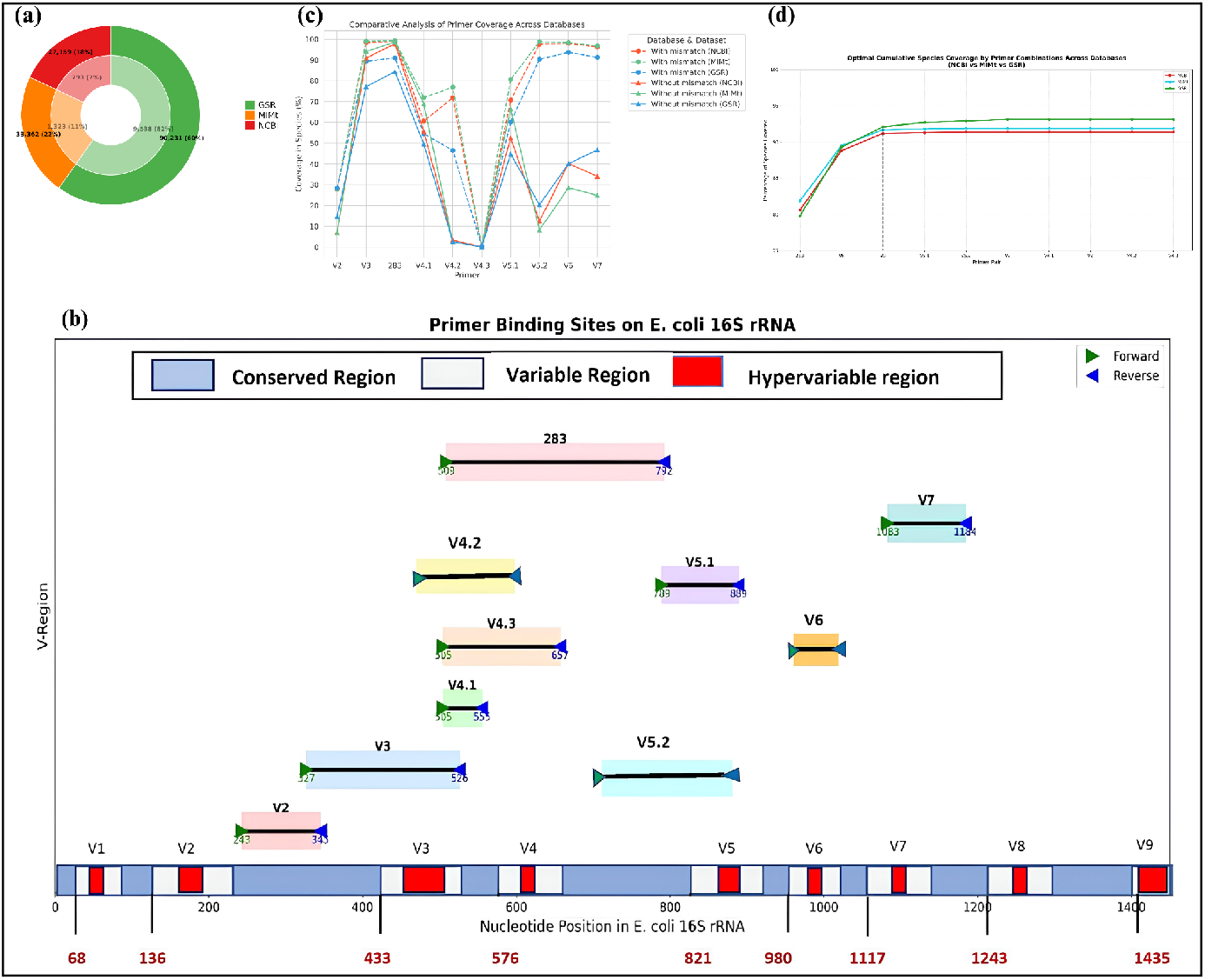
(a)Distriution of organisms in the available datasets(Outer ring - Total, Inner ring - ESKAPE), (b)Primers and their binding sites on E. coli 16S rRNA, (c)Comparative analysis of primer coverage across databases, and (d)Cumulative species coverage by primers across databases

## Geographical Distribution

Across countries, primer-deficient strains cluster disproportionately in developing settings. Pakistan (3.61%), Iran (3.00%), Argentina (2.86%), Egypt (2.50%), and India (2.74%) exhibit high proportional contributions, despite modest absolute counts (1,278 for India; <~200 for others). In contrast, developed countries with the largest counts—USA (1501) and China (1669)—exhibit lower percentages (1.26% and 1.6%, respectively), as do Germany (1.23%) and France (0.40%). Belgium (4.02%) is the lone developed-country outlier at low count. This enrichment of missing-primer isolates in developing countries, despite lower sampling depth, supports the hypothesis that reliance on common 16S primers systematically underrepresents clinically relevant strains from these regions(Figure 2).

**Figure 2:**
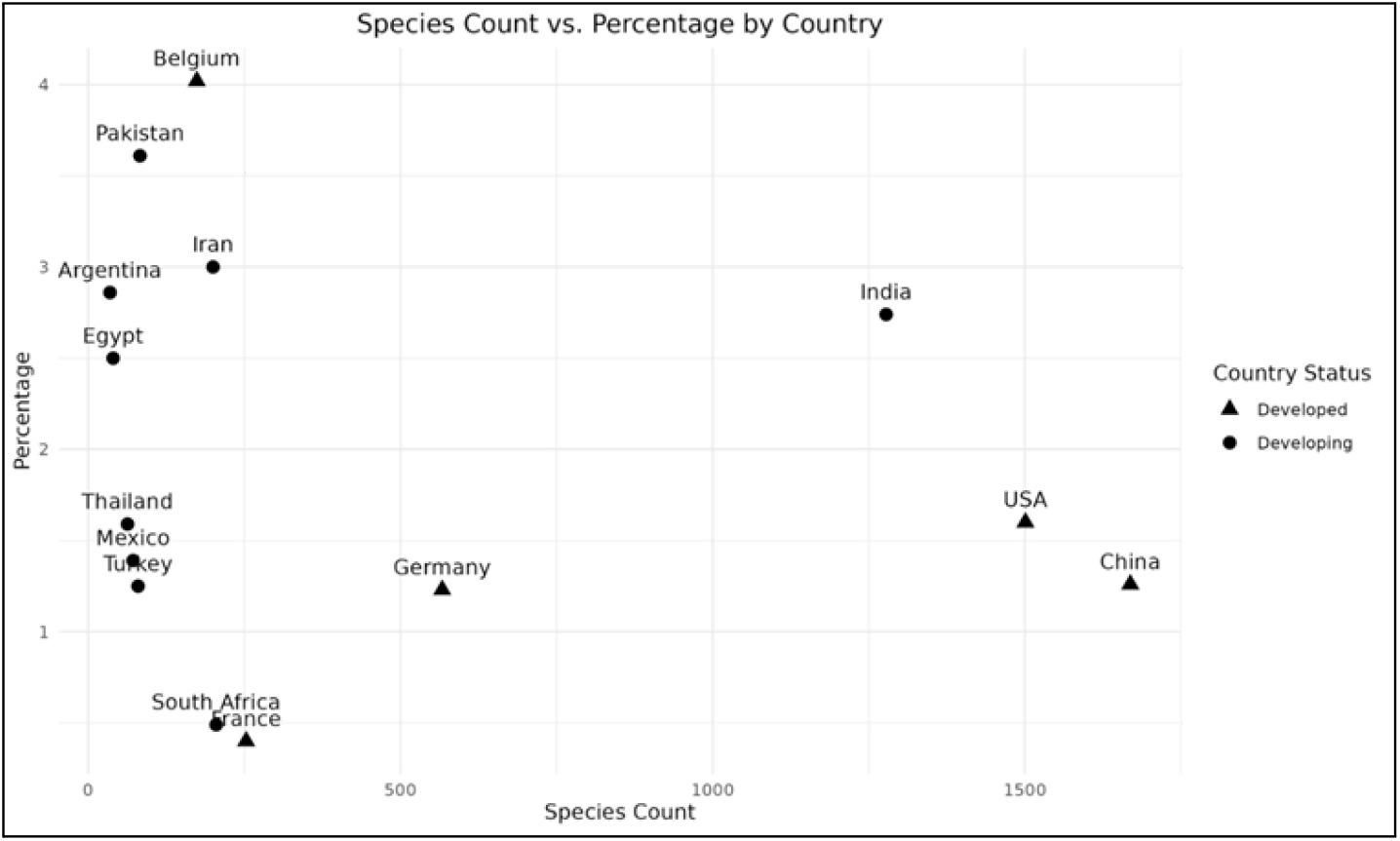
Percentage of species with suboptimal primer presence in different countries

## Discussion and Conclusion

Our results highlight a key vulnerability in the prevailing reliance on single-primer 16S rRNA sequencing for pathogen surveillance, and the geographic patterns we observed compound this issue.

A multiplexed primer approach offers a promising path forward, as it enhances species capture by spanning multiple hypervariable regions and reduces the likelihood that strains will be missed due to primer incompatibility.

Encouragingly, LMICs are beginning to take active steps to overcome these limitations. The “One Day One Genome” project, launched in 2024 by India’s Department of Biotechnology (DBT) and Biotechnology Research and Innovation Council (BRIC), aims to sequence and release a fully annotated bacterial genome from India each day. Such initiatives demonstrate both technical capacity and commitment toward building more representative genomic resources.

This study has limitations. Metadata on isolate provenance was incomplete, and some older database entries may be affected by sequencing artefacts. Moreover, in-silico binding predictions cannot fully replicate PCR efficiency under laboratory conditions. Taken together, the systematic primer biases and geographic skews underscore the urgent need for collaborative international efforts to revise and standardise primer sets, ensuring that diagnostic and surveillance pipelines are inclusive of pathogens across all global settings.

## Funding

The authors declare that no external funding was received for this study. The work was carried out using institutional and/or personal resources.

## Conflicts of Interest

The authors declare no conflicts of interest.

## Consent for publication

All authors approve this manuscript for publication.

